# Chromatin folding variability across single-cells results from state degeneracy in phase-separation

**DOI:** 10.1101/2020.05.16.099275

**Authors:** Mattia Conte, Luca Fiorillo, Simona Bianco, Andrea M. Chiariello, Andrea Esposito, Mario Nicodemi

## Abstract

Chromosome spatial organization controls functional interactions between genes and regulators, yet the molecular and physical mechanisms underlying folding at the single DNA molecule level remain to be understood. Here we employ models of polymer physics to investigate the conformations of two 2Mb-wide DNA loci in human HCT116 and IMR90 wild-type and cohesin depleted cells. Model predictions on the 3D structure of single-molecules are consistently validated against super-resolution single-cell imaging data, providing evidence that the architecture of the studied loci is controlled by a thermodynamics mechanism of *polymer phase separation* whereby chromatin self-assembles in segregated globules. The process is driven by interactions between distinct types of cognate binding sites, correlating each with a different combination of chromatin factors, including CTCF, cohesin and histone marks. The intrinsic thermodynamics degeneracy of conformations results in a broad structural and time variability of single-molecules, reflected in their varying TAD-like contact patterns. Globules breathe in time, inducing stochastic unspecific interactions, yet they produce stable, compact environments where specific contacts become highly favored between regions enriched for cognate binding sites, albeit characterized by weak biochemical affinities. Cohesin depletion tends to reverse globule phase separation into a coil, randomly folded state, resulting in much more variable contacts across single-molecules, hence erasing population-averaged patterns. Overall, globule phase separation appears to be a robust, reversible mechanism of chromatin organization, where stochasticity and specificity coexist.

## INTRODUCTION

In the cell nucleus, chromosomes are folded into a complex 3-dimensional (3D) architecture^1–5^ including a hierarchy of interactions, from *loops*^*6*^ and TADs^7,8^ to, above the megabase scale, metaTADs^9^ and A/B compartments^10^ as revealed by population-averaged contact maps^6,10–12^. Such an organization serves important functional purposes as genes and enhancers have to form specific physical contacts to regulate transcription. TADs, for instance, are thought to act as insulating structures, spatially confining the activity of enhancers to their proper targets^2,3,5^.

Different molecular factors and mechanisms have been involved in the 3D organization of chromatin. CTCF binding sites and cohesin have been proposed to shape *loops* and TADs^6^, for example via the cohesin/CTCF based loop-extrusion model^13–15^. However, while acute depletion of CTCF or cohesin leads to *loop* loss in bulk Hi-C data, signals persist at the compartment level and finer contact patterns remain within former *loops* or TADs^16–18^. Compartments A and B are known to correlate to different transcriptional states^10^, and homotypic interactions between active and poised gene promoters, linked respectively to Pol-II-S2p and PRC2, have been observed at the Mb scale and traced back to phase separation mechanisms^19–21^. Indeed, phase separation has emerged as a paradigm of cell organization^22^ and of transcriptional control^23^, as combinations of Pol-II with transcription factors and coactivators, such as Mediator, appear to form condensates^24–26^, or more fleeting interactions^27^, linked to gene regulation^23,28–30^. Yet, it remains unclear how those mechanisms act and combine to shape chromatin architecture.

Single-cell Hi-C experiments, for example, have highlighted the stochastic nature of TADs and the strong variability of their contacts^31–34^. Recent super-resolution imaging approaches have shown that TAD-like structures are present in single-cells with chromatin folded in globular 3D conformations, but they broadly vary from cell to cell^35–39^. In particular, TAD boundaries were discovered to occur with nonzero probability at all genomic positions and to have enrichments associated to only a subset of the CTCF sites in the considered regions^37^. In addition, cohesin depletion was found to leave contact patterns at the TAD-scale intact in single cells, albeit domain boundaries become equally likely to locate at any genomic position, hence abolishing TADs at the population-average level. That hinted that chromatin contacts can arise from mechanisms distinct from the loop-extrusion^37^.

Those diverse results raise questions on the nature and origin of contact patterns in single DNA molecules. Are there other folding mechanisms beyond loop-extrusion? How does phase separation act? If interactions are stochastic, how is specificity controlled? What is the origin of structural variability across cells and in time? To attack those questions, here we use a chromatin model from polymer physics to derive predictions about DNA single-molecule 3D structures that we compare with super-resolution imaging data in single-cells^37^. In particular, we investigate two 2Mb wide DNA regions in human HCT116 and IMR90 cells, where bulk Hi-C^6,16^ and single-cell imaging^37^ data are available. To reconstruct chromatin 3D conformations different computational methods^40–43^ and polymer models have been developed^13,14,15,19,20,44-52^. In this work, we focus on the textbook scenario where contacts between distal DNA binding sites are established by diffusing cognate binding factors, as described by the *Strings&Binders* (SBS) polymer physics model of chromatin^19,20,47^ (**Fig. 1a**). By machine learning from only Hi-C data^6,16^, we infer the genomic location of the putative binding sites of the SBS polymer model of the loci of interest, which are shown to correlate with specific combinations of known chromatin organizing factors. Next, by Molecular Dynamics simulations we derive a thermodynamics ensemble of single-molecule 3D structures of those loci.

**Figure 1.**
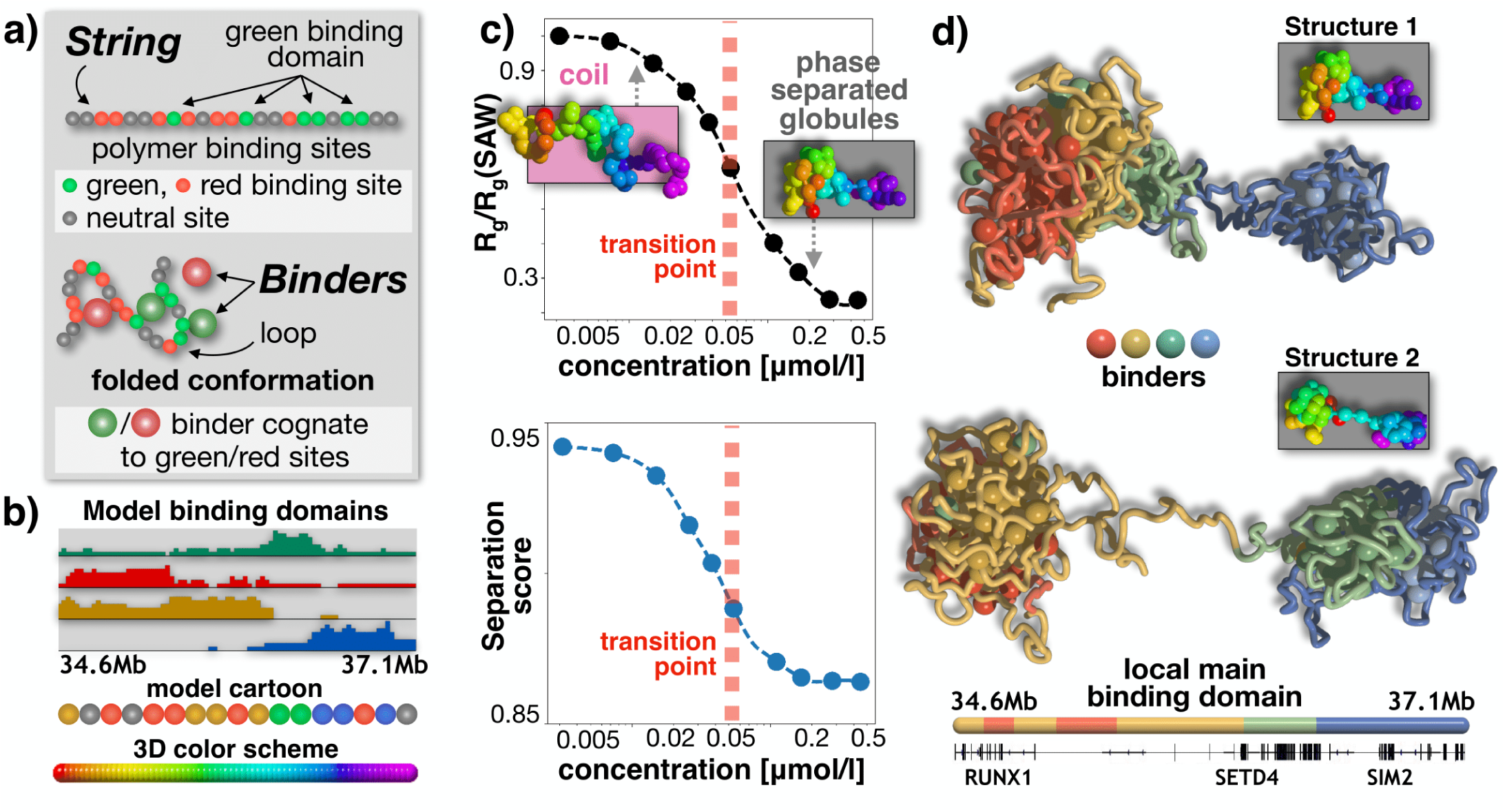
The model phase transition from a coil to a globule phase separated state. **a)** Cartoon of the *Strings&Binders* (SBS) polymer model of chromatin showing the specific binding sites along the chain (top) and a 3D conformation of the system folded by the action of cognate binders (bottom). **b)** The SBS model of the studied chr21:34.6-37.1Mb locus in human HCT116 cells has four types of binding sites, forming four *binding domains* each represented with a different color. Their genomic location and abundance are shown. A cartoon is sketched (bottom) of the model polymer chain and the color scheme used in the 3D representations in panel c). **c)** Upon increasing the binder concentration, the model has a phase transition from a *coil* to a *globule phase separated* state, a more condensed structure made of partially separated globules, as signalled by a sharp decrease of the system *order parameters*, respectively the equilibrium gyration radius (top) and average separation score (bottom). **d)** The intrinsic degeneracy of the globule phase separated state, enhanced by the overlapping genomic organization of the binding domains, corresponds to a variety of 3D conformations: two structures are shown where, for example, the green binding domain collapses respectively onto the brown and the blue domain.

As dictated by polymer physics^53^, we find that the model 3D conformations fall in two main folding classes corresponding to its thermodynamics phases, the coil, i.e., randomly folded, and the globule state, where distinct globules self-assemble along the chain by the interactions of cognate binding sites. According to the concentration or affinity of binders, the system switches from one to the other state via a phase transition mechanism of *polymer phase separation*. We show that those 3D structures recapitulate bulk Hi-C data and we validate model predictions on single-molecule 3D conformations against independent imaging single-cell data in both wild-type and cohesin depleted cells^37^. The consistent agreement provides evidence that, in the studied loci, chromatin folding is explained at the single-molecule level by such a thermodynamics mechanism, different from loop-extrusion. In particular, in the model of wild-type cells we find that the loci fold mostly in globule conformations, whose inherent thermodynamics degeneracy manifests in the broad variability of TAD-like domains across single-molecules. We also explore the time dynamics of chromatin structure at the single molecule level. Globule formation produces dynamic, yet stable local compact environments highly favoring close contacts between sites enriched for cognate binding sites, within and, less frequently, across globules. That exemplifies how stochasticity of DNA interactions can coexist with contact specificity. Acute cohesin depletion reverses phase separation into the coil state in the majority of cells, producing much more variable and transient contact patterns.

## RESULTS

### Model phase transition to the globule phase separated state

We focused, first, on modelling a 2.5Mb DNA region (chr21:34.6Mb-37.1Mb) in human HCT116 cells. The SBS is a simplified, coarse-grained model where a chromatin filament is represented as a self-avoiding chain of beads and along the chain are located specific binding sites for cognate, diffusing molecular binders^19,20,47^ (**Fig. 1a**), as well as unspecific binding sites (**Methods**). To check that our general conclusions are robust, as expected from Statistical Mechanics^53^, in our study we explored a spectrum of specific and unspecific affinities between binders and binding sites in the weak biochemical energy range, respectively from 3.1K_B_T to 8.0K_B_T (for simplicity equal across the different types) and from 0K_B_T to 2.7K_B_T (**Methods**).

To infer the genomic location and the types of the putative binding sites of the SBS polymer model of the locus, we developed a machine learning procedure (**Methods** and **Supplementary Fig. 1**) based on the PRISMR approach^50^, which employs as input only bulk Hi-C data^16^, with no use of epigenetic tracks to avoid biases towards a subset of factors. The procedure returns four distinct types of specific binding sites (visually represented by different colours, **Fig. 1b**), each defining a binding domain. After setting the affinities, the system is investigated at different binder concentrations (equal for all types), from 0 to 0.5μmole/litre, by Molecular Dynamics (MD) simulations to derive, for each different concentration, a thermodynamic ensemble of single-molecule 3D conformations of the model of the locus.

Upon increasing the binder concentration, we find that at a characteristic threshold (**Fig. 1c**) the polymer undergoes a thermodynamics phase transition from a *coil* to a *globule phase separated* state^53^, corresponding to a sharp conformational rearrangement. In our HCT116 main case study, the threshold concentration is about 50nmole/litre (**Fig. 1c**) and, more generally, for the explored weak biochemical affinities it falls in the fractions of μmole/litre range^47^, values compatible with transcription factor concentrations (**Methods**). As known in block-copolymers^20,54,55^, in the coil state entropic forces keep the polymer in randomly folded conformations, while in the phase separated state attractive forces thermodynamically prevail and the different binding domains self-assemble by action of (and along with) their cognate binders in more compact and partially separated globules, as signalled respectively by a sharp drop in the gyration radius and separation score, the *order parameters* of the system (as well as in its binding energy, **Supplementary Fig. 2**, and **Methods**). Differently from usual linear block-copolymers, though, the separation of the globules is only partial because of the overlapping genomic distribution of the underlying binding sites that increases the degeneracy of the system microstates, which can fold in a multiplicity of 3D conformations (**Fig. 1d, Methods**). The self-assembly of globules is guided by the nontrivial genomic arrangement of the four binding domains of the model that are enriched each in a distinct, successive genomic region and hence form the polymer core globules, which result into the main TAD-like structures of the median distance map of the model (**Fig. 2a, Methods**).

**Figure 2.**
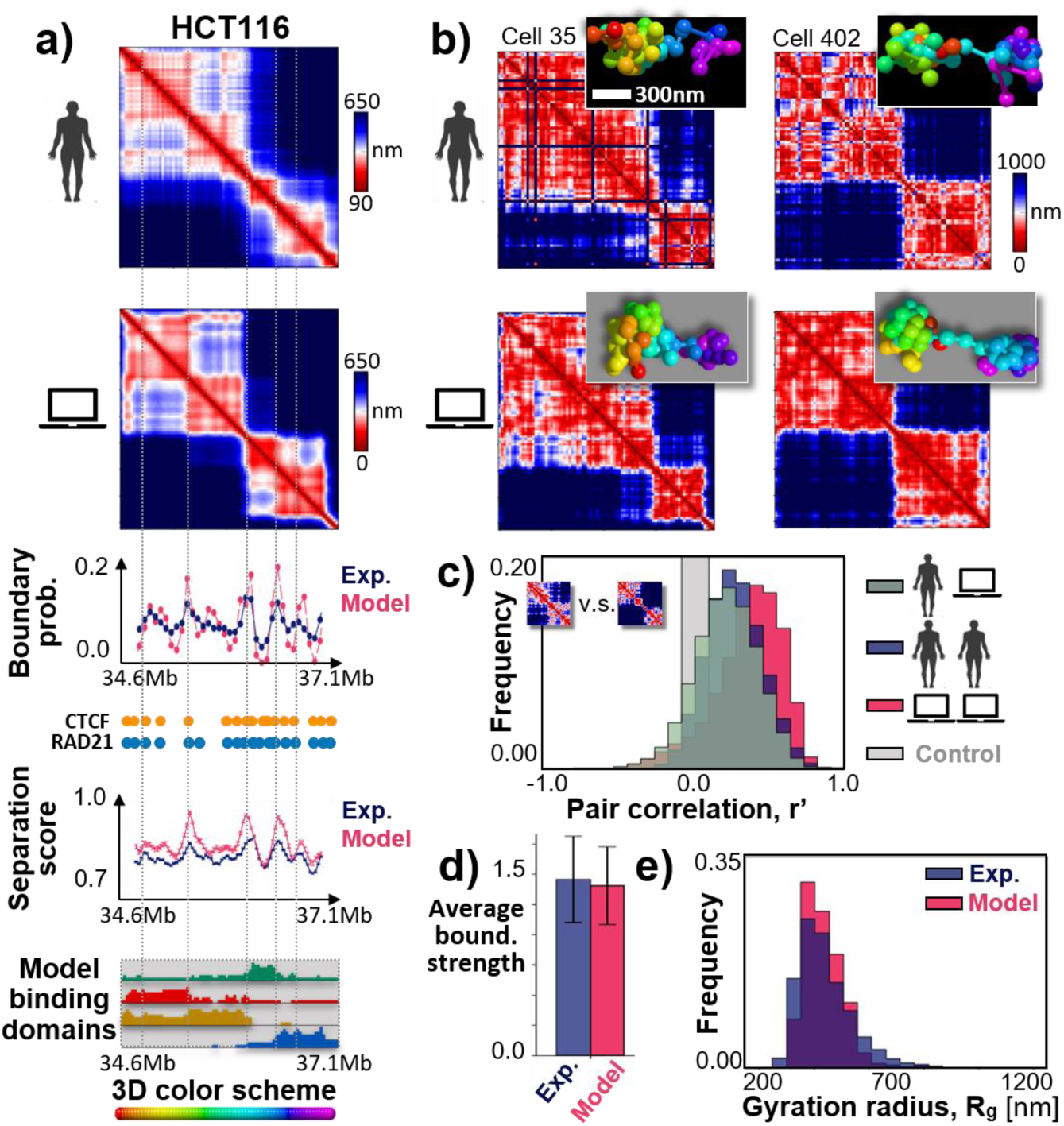
Phase separation degeneracy explains variability of single-molecule conformations. **a)** In the considered 2.5Mb wide locus chr21:34.6Mb-37.1Mb of human HCT116 cells, the model median distance matrix in the globule separated state compares well against imaging data^37^ (top, the Pearson and genomic-distance corrected correlations are, respectively, r=0.95 and r’=0.84). The probability of a domain boundary in 3D conformations along the locus (middle, model-experiment correlation r=0.79) and the corresponding separation score (model-experiment r=0.85) also match. The location of ChIP-seq CTCF (orange circles) and cohesin (RAD21, blue circles) sites^57^, and the intensity of the model four types of specific binding sites along the locus (bottom, as in Fig.1b) are also shown. The vertical dotted lines are drawn to help comparing the panels. **b)** Consistently, an all-against-all comparison of single-cell imaged^37^ (top) and model predicted (bottom) 3D structures by the RMSD method shows that all imaged conformations statistically map onto model single molecules in the globule state (bottom, see text and Suppl. Fig.s 5b, 6). The varying TAD-like domains result from the intrinsic conformational degeneracy of such a thermodynamic state. **c)** The degree of variability of single-molecules is measured by the genomic-distance corrected correlation, r’, of pairs of distance matrices. The distribution of r’ between pairs of imaged structures (blue, average r’=0.27) is statistically not distinguishable from the r’ distribution between imaged and model distance matrices (dark grey, Mann Whitney p-value=0.19). The model-model r’ distribution is in red and in light grey a control. **d)** The model and experimental average boundary strength, and **e)** the gyration radius distributions are also not distinguishable (Mann Whitney p-value=0.40). Overall, the *polymer globule phase separated* state of the model returns single molecule structures with features consistent with both single-cell imaging and bulk Hi-C (Suppl. Fig.4a) data.

To gain insights into the molecular nature of the inferred model binding sites, which are responsible of folding, we correlated their genomic positions with available epigenetic data in the same cell type^16^ (**Supplementary Fig. 3**). Interestingly, we find that each single binding type (colour) has statistically significant Pearson correlations (**Methods**) with a specific combination of known architecture organizing factors. The first putative binding domain (green, in **Fig. 1b**) correlates mainly with the CTCF/Smc1 (Cohesin) system, the second one (red) with active marks (e.g., H3K27ac and transcription factors) and less with Smc1, the third (brown) with repressive marks (e.g., H3K27me3), whereas the fourth (blue) with H4K16ac and specific transcription factors.

Summarising, our polymer model undergoes a phase transition from a coil to a phase separated globular state as the number of binders (or affinity strength) grows above a threshold point. For a given binder concentration, the system can fold in a variety of 3D conformations, not just in a unique, *naïve* structure. As dictated by polymer physics^53^, however, the system 3D conformations fall in two main folding classes corresponding to its thermodynamics phases, the coil and the globule separated states. Folding is controlled by the system binding sites and cognate binders, each type correlated with a different combination of chromatin architecture factors.

### Model validation against independent imaging distance data

To check that the model derived 3D structures recapitulate the Hi-C data used to infer its putative binding sites, we computed the average contact matrix in the two thermodynamic phases. While in the coil state the contact matrix is structureless, in the globular state it exhibits a pattern of TADs and sub-TADs similar those in Hi-C data (**Supplementary Fig. 4a**), as highlighted by the high Pearson, r=0.88, and genomic distance corrected Pearson correlation coefficient, r’=0.68, between model and Hi-C contact data (**Methods**).

In a first validation of our model and of its Hi-C inferred putative binding sites, we also compared its predictions about the locus median distance matrix in the globular state against independent super-resolution imaging data^37^ (**Fig. 2a**) and found that they have a Pearson, r=0.95, and distance-corrected correlation, r’=0.84, even higher than correlations with Hi-C data. Hence, the basic physics ingredients of our polymer model and its inferred binding sites are sufficient to recapitulate bulk Hi-C and independent imaging data.

Next, to demonstrate that our model provides a *bona-fide* representation of chromatin conformations in single-cells, we performed an *all-against-all comparison* between its predicted single-molecule 3D structures and single-cell 3D structures from imaging data^37^ (**Fig. 2b**). By use of a method^33^ that finds the optimal rotation between two centered 3D structures to minimize the mean squared deviation (RMSD) of their coordinates (**Supplementary Fig. 5a**), each experimental 3D structure was univocally associated to a corresponding model 3D structure by searching for the least RMSD (**Supplementary Fig. 6a** and **Methods**). Consistent with the results on average contact and distance matrices, in the HCT116 case we find that all experimental structures map onto model conformations in the thermodynamics globule state (**Supplementary Fig. 5b**). To test the significance of the association, we compared the RMSD distribution of the experiment-model optimal matches to the RMSD distribution of pairwise comparisons between experimental structures (null model): the two distributions are statistically different (Mann-Whitney test p-value=0) with only 2% of entries of the former falling above the first quartile of the latter (**Supplementary Fig. 6b**). Additionally, we find that each model globule conformation is significantly associated to at least one experimental structure, showing that the model well represents the experimental ensemble (**Methods**).

### Degeneracy in phase separation explains variability of single-molecule conformations

To further validate our model, we compared the architectural features of its predicted single-molecule 3D conformations against single-cell 3D structures from imaging^37^ (**Fig. 2b**). In single cell experiments, the locus folds in spatially segregated globules, as highlighted by the separation score as a function of the genomic coordinate (**Fig. 2a**), which produce the TAD and sub-TAD-like domains of the distance matrix. However, the 3D structures are broadly varying across single-cells, and TAD boundaries are found to be spread along the entire locus (see the boundary probability in **Fig. 2a**). We aimed to test whether the model ensemble of single-molecule conformations has features similar to those found in single-cell experiments and whether it has a similar variability (**Fig. 2b**).

First, we found that: *i)* the model derived TAD-like boundary probability and, *ii)*, separation score along the locus are very similar to the experimental ones (respectively r=0.79 and r=0.85, **Fig. 2a** and **Methods**); *iii)*, the average boundary strengths are similar (**Fig. 2d**); *iv)* the average boundary probabilities and, *v)*, the boundary strength distributions are similar too (**Supplementary Fig**.**s 7a, b**), albeit there are no free parameters in all those comparisons. Additionally, the gyration radius distributions of the model and experiment are also found to be statistically not distinguishable from each other (Mann Whitney p-value=0.40, **Fig. 2e**). Conversely, a control block-copolymer model with four non-intertwining binding domains designed specifically to reproduce the main TAD-like structures visible in bulk Hi-C data, which has also a coil-to-globule transition, was found to poorly reflect the complexity of the observed contact patterns (**Supplementary Figure 8** and **Methods**).

Second, to quantify the variability of experimental single-cell 3D structures, we measured the distance-corrected correlation, r’, between pairs of single-cell distance matrices, and found that it has a broad distribution with an average correlation r’=0.27 (**Fig. 2c** and **Methods**, similar results are found for the Pearson correlation, r). We found that the model-model r’ distance correlation has a similar distribution and, additionally, the distribution of correlations between model and experimental single-molecule distance matrices (average r’=0.22) is not statistically distinguishable from the one between experiments (**Fig. 2c**, Mann Whitney p-value=0.19, **Methods**).

Those results show that the features of the 3D structures predicted by our model are similar to those observed in single-cell experiments, to the point that single-molecules from the model are statistically indistinguishable from experimental single-cell structures. Finally, we implemented our modelling and all the above analyses in another 2Mb locus (chr21:28Mb-30Mb) investigated in human IMR90 cells by super-resolution imaging experiments^37^ and found analogous results (**Supplementary Fig**.**s 3, 4, 7, 9, 10** and **Methods**).

The overall agreement between single-cell imaging data and the independently derived model conformations supports the view whereby, in the studied HCT116 and IMR90 loci, chromatin folding is explained at the single-cell level by a thermodynamics mechanism of globule phase separation, driven by the interactions of a few different types of binding sites, non-trivially arranged along the genome and each associated to specific combinations of chromatin organizing factors, including, but not limited to CTCF (**Fig. 2a**). Within that framework, the broad variability of single-molecule 3D globular structures, reflected in the varying locations of TAD-like domain boundaries, naturally results from the inherent folding degeneracy of the phase separated conformations, enhanced by the overlapping genomic organization of the different binding domains. Whereas CTCF sites are distributed over the entire locus, the boundary preferential positions correspond to the location of the edges between binding domains (**Fig. 2a**) as they are prone to fold in separated globules.

### Cohesin depletion reverses phase separation

To investigate how acute cohesin depletion impacts single-molecule chromatin conformations, we considered the same locus in HCT116 Auxin treated cells (HCT116+Auxin)^37^. We inferred the new SBS polymer binding sites, as before, from Hi-C data in HCT116+Auxin cells^16^ and derived by MD the model 3D conformations to be compared with imaging data in the new cells^37^. Interestingly, in this case our approach finds only three types of specific binding sites in the locus (**Fig. 3a**). The domain strongly correlated with cohesin in wild-type (WT) HCT116 cells (green, **Fig. 2a**) disappears, whereas the other WT domains are overall maintained at their genomic locations, although weakened and shrunk, and their epigenetic signatures partially preserved (**Fig. 3a** and **Supplementary Fig. 3b**). We find that the new polymer model also undergoes a phase transition from a coil to a globule phase separated state, yet at around 400nmole/litre if the same affinities of the HCT116 case study model are used (**Supplementary Fig. 2** and **Methods**).

**Figure 3.**
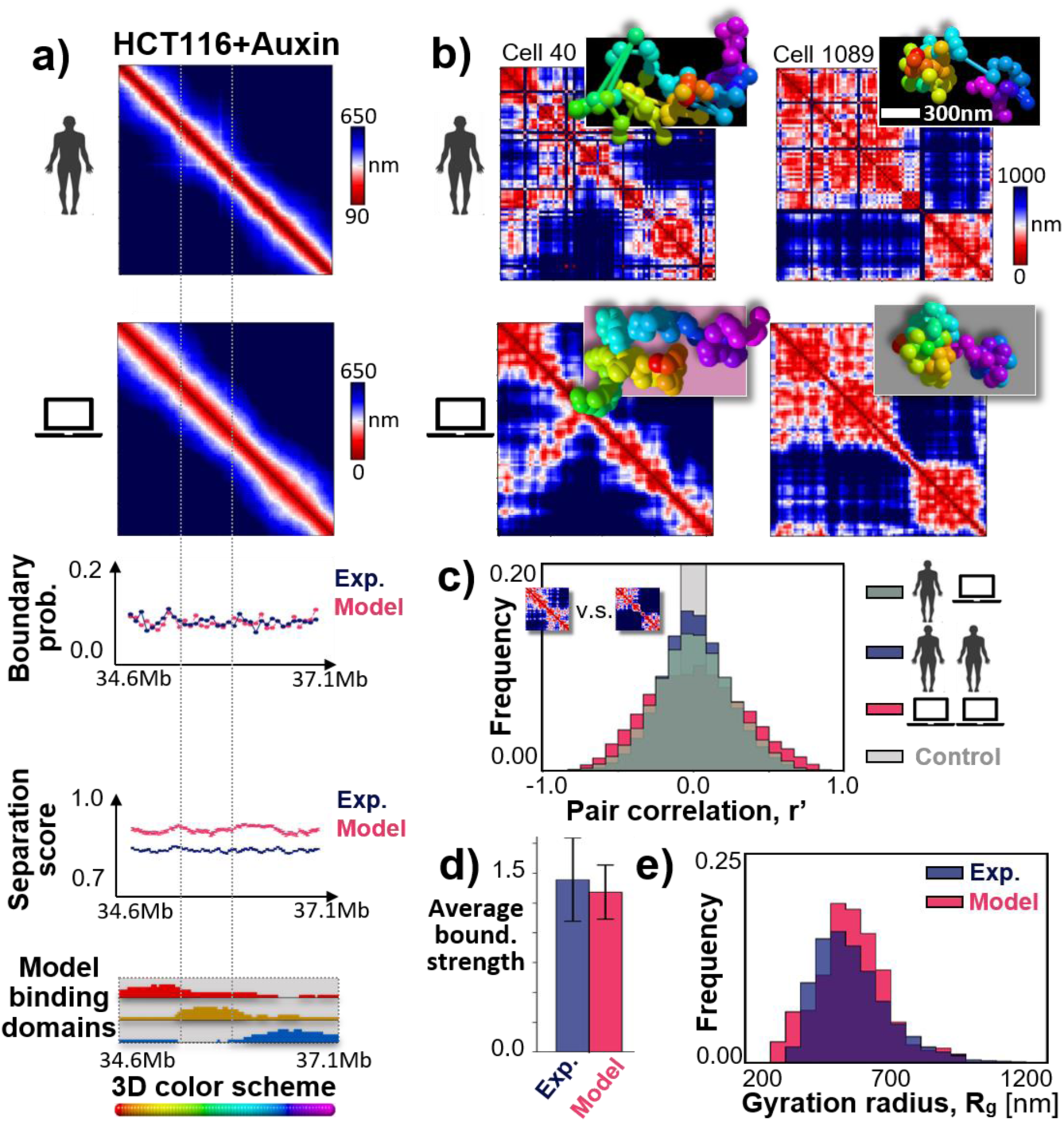
Cohesin depletion tends to reverse phase separation. **a)** Top: In cohesin depleted HCT116 cells treated with Auxin (HCT116+Auxin), the model predicted median distance matrix of the considered locus chr21:34.6Mb-37.1Mb also compares well against independent imaging data^37^; the Pearson and genomic-distance corrected correlations are, respectively, r=0.96 and r’=0.57. A mixture of model 3D structures is required, however, 80% in the coil and 20% in the globule state. Middle: The flat domain boundary probability and separation score reflect the absence of TAD-like structures in the median matrices. Bottom: The model of the locus in cohesin depleted cells has three binding domains; their different colors are assigned by their genomic overlap with the wild-type domains of Fig. 2a. **b)** Consistently, the RMSD based all-against-all comparison of single-cell^37^ (top) and model predicted (bottom) 3D structures shows that imaged conformations correspond to model structures belonging 80% to the coil (bottom left) and 20% to the globule state (bottom right, see text and Suppl. Fig.s 5c, 11). The distance matrix of single molecules has non-trivial patterns in both states, but in the coil state (bottom left) contacts originate from random collisions rather than stable phase separated globule domains (bottom right). **c)** The distribution of genomic-distance corrected correlations between distance matrices from single-cells (blue) is broader than in wild-type and its average is r’=0.0, highlighting a higher variability; it is statistically not distinguishable from the correlations between imaged and model distance matrices (dark grey, Mann Whitney p-value=0.48). **d)** In model and experiment the average boundary strength is similar, and similar to wild-type (Fig. 2d), and **e)** the gyration radius distributions are not distinguishable too (Mann Whitney p-value=0.10) and have a higher average value than wild-type (540nm v.s. 440nm of Fig. 2e). The model-experiment agreement points out that cohesin depletion reverses phase separation in most cells as their corresponding model single-molecule structures are mainly in the coil rather than in the globule state, contrary to HCT116 (Fig.2).

The Hi-C map of the cohesin depleted locus lacks the wild-type TAD-like structures and retains only a faint pattern of interactions^16^. The model recapitulates well those data too (r=0.93, r’=0.33, **Supplementary Fig. 4b**), but we find that a mixture of 3D structures is required, composed 80% of single-molecule 3D conformations in the coil and 20% in the globule phase separated thermodynamics state. Consistently, in the HCT116+Auxin case by the least RMSD method we find that 80% experimental structures from independent imaging data^37^ (**Fig. 3b**) map onto model conformations in the coil and 20% in the globule state (**Supplementary Fig. 5c**) in a statistically significant association (**Supplementary Fig. 11**). Again, the comparison of our mixture model prediction on the median distance matrix against the independent imaging data^37^ gives high correlations (r=0.96, r’=0.57, **Fig. 3a**).

Upon cohesin depletion, although the population-averaged distance map is as featureless as the Hi-C map, in single-cell imaging data contact patterns persist, including TAD-like structures in some instances (**Fig. 3b**). The domain boundary strength and the average number of boundaries are similar to WT^37^. However, the imaged single-cell 3D conformations have a higher variability than WT ones: the average distance-corrected correlation, r’, between pairs of distance matrices is r’=0.0 and its distribution is broader (**Fig. 3c**). The model single-molecule conformations have also a high variability and resemble the experimental structures (**Fig. 3b**). Again, they have an r’ correlation distribution with imaged distance matrices (and with each other, average respectively r’=0.0 and r’=0.0) statistically similar to the one between experiment pairs (Mann Whitney p-value=0.48, **Fig. 3c**). The 3D conformations of the model mixture include globular states as in wild-type (**Fig. 3b** right), but 80% of single-molecules are in the coil state (**Fig. 3b** left) whose contact patterns reflect transient, random chromatin collisions rather than more stably folded contacts as in wild-type (see time dynamics section below). Consistent with such a picture, the average separation score is flat along the locus in both model and experiment (**Fig. 3a**). The model domain boundary probability along the locus is also as flat as the experimental one (**Fig. 3a**) with a similar average boundary strength (**Fig. 3d**); and similar are the average boundary probability and the boundary strength distribution (**Supplementary Fig**.**s 7c, d**), as much as the gyration radius distribution (Mann Whitney p-value=0.10 **Fig. 3e**), whose average value is 23% larger than in the wild-type case (540nm v.s. 440nm) showing that the locus is more open.

The overall agreement between model and independent microscopy data in the HCT116+Auxin case depicts a scenario where, consistent with the known role of cohesin as a key architecture organizing factor, cohesin depletion reverses chromatin globule phase separation to the coil thermodynamics state in single-cells, whose diverse contact patterns originate mainly from random chromatin collisions rather than from phase separated domains.

### Single-molecule time dynamics

Next, we investigated how the spatial conformations of single DNA molecules change in time and how specific patterns of contact or insulation are established, which can be uniquely achieved within our model. In the steady-state, the 3D structure of a single-molecule varies and breathes under thermal fluctuations in both the coil and phase separated states, but important differences mark the two phases (**Fig. 4**).

**Figure 4.**
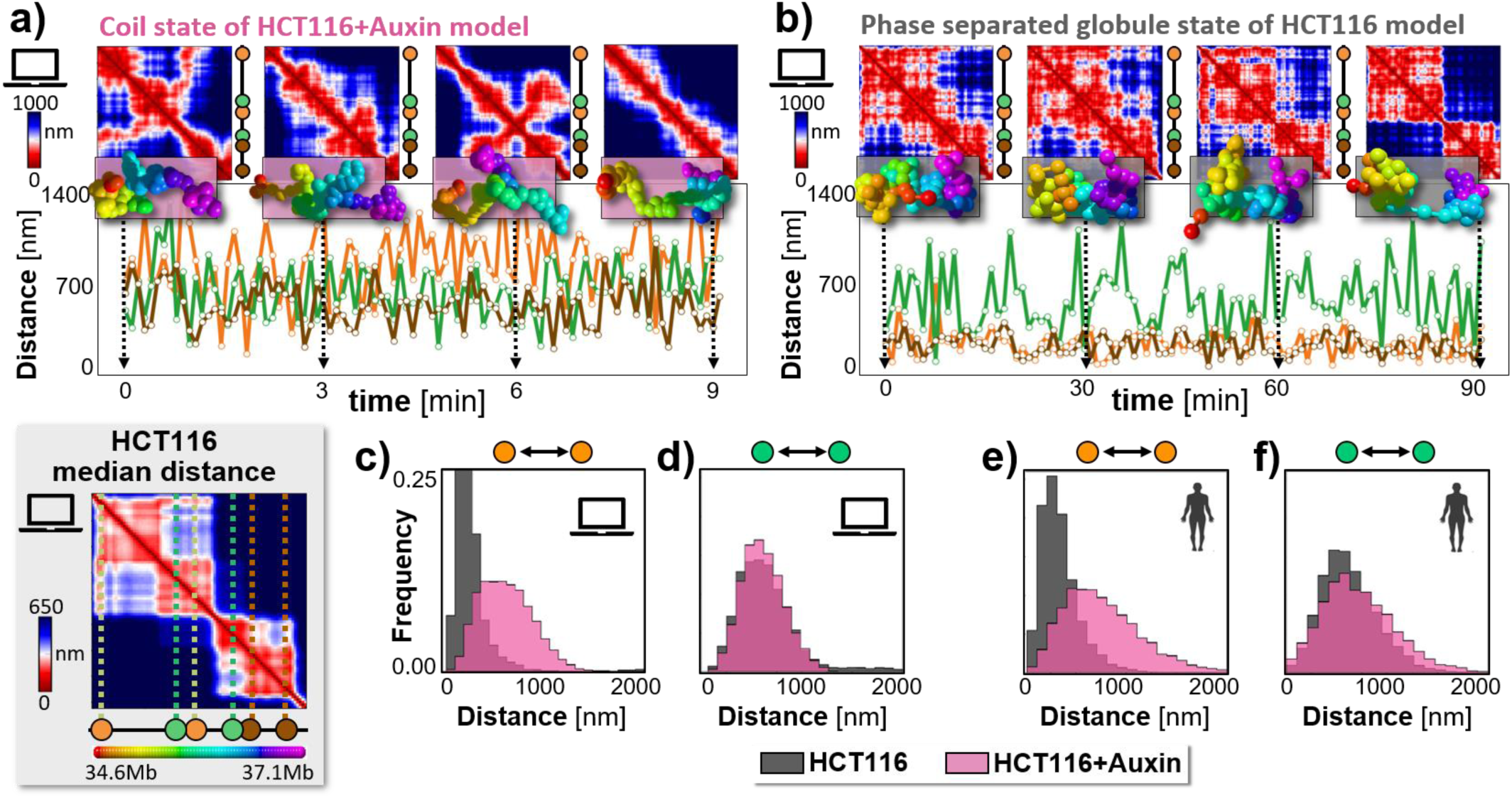
The time dynamics of single molecules illustrates how globules establish specific contacts and boundaries. The steady-state time behavior is shown of a single-molecule and its distance matrix in the model of the chr21:34.6Mb-37.1Mb locus in cohesin depleted (HCT116+Auxin coil state, panel **a)**) and wild-type cells (HCT116 globule state, panel **b)**). The relative distances are also plotted of: *i)* a pair of sites (orange), 1.2Mb apart, in different subTADs, having a strong point-wise (*loop*) interaction in WT HCT116; *ii)* a pair of 0.6Mb distant sites (green) with a TAD boundary in between; *iii)* a pair of sites (brown), almost 0.6Mb apart within the same subTAD. **a)** In the coil state, the distance time tracks of those sites have all wide fluctuations, as contact patterns are fleeting. **b)** In the globule phase separated state, the interaction pattern shows that globules vary, but persist in time (see text). Hence, within their stochastic environment, close, specific contacts are enhanced between site pairs sharing abundant cognate binding sites (orange and brown, Supplementary Fig. 13), whereas pairs in different globules remain insulated (green). **c)** The orange pair is on average much closer (Supplementary Table I) and its distance distribution much narrower in the HCT116 than in the HCT116+Auxin model, **d)** whereas the green pair behaves similarly in both. **e, f)** The model distributions are close to the corresponding experimental ones^37^ in HCT116 and in HCT116+Auxin cells, consistent with the view that chromatin folds in different thermodynamics states in WT and cohesin depleted cells.

In the coil state, the contacts visible in the distance matrix of a single molecule have a highly transient nature and their pattern fully changes in time (**Fig. 4a**, HCT116+Auxin model), as signaled by the average value of the r’ correlation between different time points that approaches zero for large time separations (**Supplementary Fig. 12a**), consistent with the zero average correlation between different replicates discussed before. In the phase separated state, the 3D structure also varies in time, but the long-time average r’ correlation remains well above zero (in the HCT116 model r’ plateaus to 0.39, **Supplementary Fig. 12b**), showing that the folded globules change, but persist in time (**Fig. 4b**), again consistent with the average non-zero correlation between replicates. The conformation average decay time (i.e., the time for correlations to plateau) is almost one order of magnitude larger in the globule state than in the coil state; its scale can be roughly guessed by using estimates of the viscosity of the nuclear medium reported in the literature^45,56^: for example, it results to be 9sec and 60sec respectively in the coil state of the HCT116+Auxin and in the phase separated state of the HCT116 model (**Supplementary Fig. 12** and **Methods**).

Finally, we explored how domain boundaries and specific contact *loop*s are established at the single-molecule level, in the face of a varying environment, by the formation of globules. To that aim, we investigated the relative distances of a particular set of sites: *i)* a pair of sites (orange, **Fig. 4**) having in HCT116 cells a strong point-wise (*loop*) interaction in bulk Hi-C data, albeit located 1.2Mb apart from each other in different subTADs; *ii)* a pair of 0.6Mb distant sites (green) with a strong TAD boundary in between; *iii)* a control pair of sites (brown), almost 0.6Mb apart, enclosed within a subTAD.

In the HCT116+Auxin model, where molecules are mostly in the coil state, the average physical distances of the green and brown pair are comparable to each other (around 620nm, **Supplementary Table I**) and the orange pair is more open (660nm) for its larger genomic separation. The distance distributions are comparatively broad and similar across the three pairs (**Fig. 4c, d** and **Supplementary Fig. 13a**). The situation drastically changes in the globule phase separated state of the model of HCT116 cells as the average distance of the orange and of the brown pairs is reduced of factor 2.5 down to around 280nm. That occurs because the orange (and brown) genomic regions are enriched with cognate binding sites, which in their globule compact environment are highly likely to be bridged hence resulting in a *loop* visible in Hi-C bulk data. Conversely, the green sites tend to become trapped each in a different globule, remaining at roughly their coil-state distance. In this way, globules form an insulating “boundary” between them. The distance distribution of the orange (and brown) pair is much narrower in the HCT116 than in HCT116+Auxin case, whereas the distribution of the green pair is similar in both (**Fig. 4c, d** and **Supplementary Fig. 13a**).

We performed an initial validation of the model time behavior by comparing the predicted distance distributions of the mentioned site pairs with single-cell imaging data, although a full test would need experiments following in real time the entire chromatin locus. Interestingly, considering the basic character of the model, its predicted distributions are comparatively close to the experimental ones, albeit there are no free parameters available in the comparison (**Fig. 4e, f** and **Supplementary Fig. 13b**). That is consistent with the above interpretation that chromatin folds in different thermodynamics states in wild-type and cohesin depleted cells. Finally, the time tracks (**Fig. 4a, b**) also clarify that the distances of all site pairs change in time subject to thermal fluctuations and, in particular, the strong point-wise *loop* interaction of the orange pair visible in the median distance matrix in HCT116 cells does not reflect a fixed-length permanent contact. Again, analogous results are found for the locus in IMR90 cells (**Methods**).

Overall, the analysis of the steady-state time dynamics shows that, while in the coil state contacts within a single molecule are fleeting and variable, in the phase separated state globules breathe and rearrange, but persist in time, as discussed in polymer physics^53^. Hence, globules can create spatially compact environments, visible as TADs and sub-TADs in Hi-C data, where specific contacts (e.g., the loops of the brown and orange pairs) are enhanced between regions sharing abundant cognate binding sites, albeit based on weak biochemical interactions. Globule boundaries also change in time, but they can efficiently separate neighboring regions along the sequence (see, e.g., the green pair), although specific contacts across proximal globules can also form (e.g., the loop of the orange pair).

## CONCLUSIONS

DNA loop-extrusion has recently emerged as an important mechanism of chromatin organization^13–15^. It envisages that a cohesin complex acts as an *active motor* extruding loops between CTCF anchor points, in a non-equilibrium process requiring energy influx to work, e.g., ATP molecule consumption. The key role of CTCF/cohesin in chromatin architecture has been confirmed, for example, by bulk Hi-C data in systems depleted for those factors^16–18^. However, in the 2Mb-wide loci in human HCT116 and IMR90 cells considered here, super-resolution single-cell imaging experiments questioned the loop-extrusion scenario, hinting that DNA interactions can arise from a distinct molecular process^37^. Here, we discussed a mechanism of chromatin folding, different from the loop-extrusion, that is based on the thermodynamics of polymer phase separation and is consistent with both Hi-C and single-cell imaging data.

Specifically, we considered a schematic polymer model of chromatin, the *Strings&Binders* model^19,20^, where contacts between distal binding sites are mediated by diffusing cognate bridging molecules (but our results also hold if DNA sites have direct physical interactions rather than mediated by binders, **Methods**). The genomic arrangements of the model putative binding sites are learned from Hi-C bulk data^6,16^ of the loci of interest, and the thermodynamics 3D conformations of the system derived from physics. Upon increasing the binder concentration, or affinity, the model undergoes a phase transition from a coil to a globule phase separated state where compact globules self-assemble by the interactions with their cognate binders. Importantly, as dictated by polymer physics^53^, the model 3D structures spontaneously fall in the conformational class corresponding to its thermodynamics phase, i.e., the coil or globule state (**Fig. 5a**). The consistent agreement between the predicted structures and independent single-cell super-resolution microscopy data^37^ provides evidence that, in the studied loci, chromatin folding is driven at the single-molecule level by such a mechanism of *polymer phase separation*.

**Figure 5.**
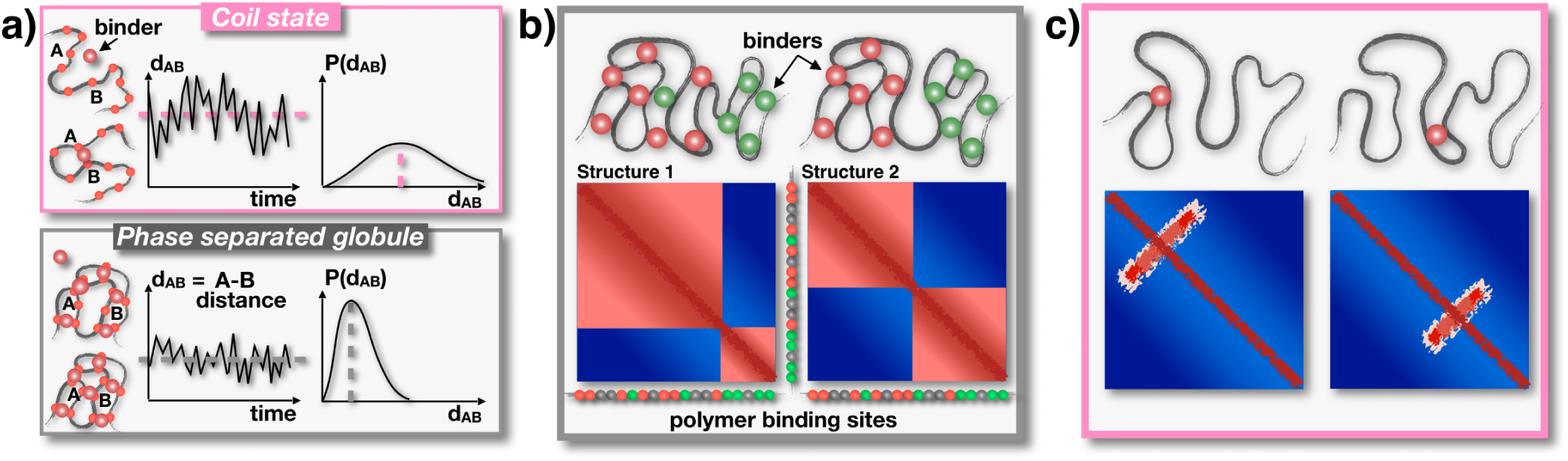
Polymer phase separation explains chromatin structure variability across single-cells. **a)** In the SBS model, the polymer folds in conformational classes associated to the thermodynamics phases of the system, e.g., the coil and the phase separated globule state, according to the abundance or affinity of binding molecules. The summary cartoon highlights that in the coil phase (top) chromatin contacts tend to be more fleeting and average distances more open than in the phase separated globule state (bottom). Those thermodynamics states recapitulate single-cell architecture data. **b)** Globule phase separation, for example, produces local compact environments, insulated one from the other, where specific contacts are highly enhanced between regions enriched for cognate binding sites. The inherent thermodynamic degeneracy of conformations results in a variety of single-molecule 3D structures, reflected in the variability of the genomic position of TAD-like patterns, consistent with imaging data of loci in human HCT116 and IMR90 cells. **c)** Cohesin depletion, as in HCT116+Auxin cells, tends to reverse globule phase separation into the coil state resulting in much more variable contacts, abolishing patterns in bulk Hi-C data.

The emerging scenario shows that in WT cells the loci fold mostly in the globule phase separated state, whose intrinsic thermodynamics degeneracy is manifested in the varying genomic positions of TAD-like patterns across single-molecules and in time (**Fig. 5b**). Population-averaged contact maps, such as Hi-C bulk data, capture ensemble averages and their TADs match the location of the globules that more frequently form. The analysis of the time dynamics of single molecules illustrates the diverse modes of action of globules in shaping spatial interactions or insulation between distal sites. While segregating neighbouring regions, they create stable, compact local environments enhancing specific contacts between sites enriched for cognate binding sites, within and less frequently across sub-TADs and TADs. That explains how the observed stochasticity of DNA interactions, typical of weak biochemical affinities, can coexist with specificity, providing a quantitative picture on how contacts, e.g., between genes and distal regulators can be controlled at the molecular level. Finally, our results are consistent with a scenario where acute cohesin depletion tends to reverse globule phase separation into the coil state in most cells, resulting in much more variable and transient contact patterns in single molecules (**Fig. 5c**), hence abolishing population-averaged TAD-like domains. We find that the model inferred binding site types have significant correlations each with a specific combination of chromatin architecture factors, rather than a single one, including CTCF, Smc1, H3K27ac or H3K27me3. That strengthens the view that the combinatorial action of different molecules, modulating each other activity, shapes the 3D architecture of the genome.

We explored a minimal model of strings and binders, but a huge diversity of microphase and phase separated structures, well beyond TAD or pled-like patterns, can be achieved by adding molecular parameters to the system^54,55^, although whether true equilibrium self-assembly can be reached in such complex systems remains to be clarified. Nevertheless, an organizational mechanism based on phase transitions has the advantage to be a robust and reversible procedure to trigger conformational changes: the system only needs, e.g., to establish an above threshold concentration of binders (or affinity), with no need of fine tuning their number (or strength)^19^. And phase transitions occur spontaneously sustained by the thermal bath. That could explain how simple cell strategies of up-regulation of genes associated to transcription factors or epigenetic modifications can reliably shape the self-assembly of chromatin architectures in the nucleus.

## ACKNOWLEDGEMENTS

M.N. acknowledges support from the NIH grant ID 1U54DK107977-01, the EU H2020 Marie Curie ITN n.813282, CINECA ISCRA ID HP10CYFPS5 and HP10CRTY8P, Einstein BIH Fellowship Award (EVF-BIH-2016-282), Regione Campania SATIN Project 2018-2020, and computer resources from INFN, CINECA, ENEA CRESCO/ENEAGRID (Ponti et al., 2014) and *Scope*/*ReCAS* at the University of Naples.

## AUTHOR CONTRIBUTIONS

M.N. designed the research project. M.C., L.F., S.B., A.M.C., A.E. developed modelling, run simulations and performed data analyses. M.N., M.C. and S.B. wrote the manuscript.

